# Commercial gene expression tests for prostate cancer prognosis provide paradoxical estimates of race-specific risk

**DOI:** 10.1101/604058

**Authors:** Jordan H. Creed, Anders E. Berglund, Robert J. Rounbehler, Shivanshu Awasthi, John L. Cleveland, Jong Y. Park, Kosj Yamoah, Travis A. Gerke

## Abstract

**Background:** Commercial gene expression signatures of prostate cancer (PCa) prognosis were developed and validated in cohorts of predominantly European American men (EAM). Limited research exists on the value of such signatures in African American men (AAM), who have poor PCa outcomes. We explored differences in gene expression between EAM and AAM for three commercially available panels recommended by the National Comprehensive Cancer Network for PCa prognosis. *Materials and Methods*: 232 EAM and 95 AAM patients provided radical prostatectomy specimens. Gene expression was quantified using Nanostring for 60 genes spanning the Oncotype DX Prostate, Prolaris, and Decipher panels. A continuous expression-based risk score was approximated for each. Differential expression, intrapanel co-expression and risk by race were assessed.

**Results and limitations:** Clinical and pathologic features were similar between AAM and EAM. Differential expression by race was observed for 48% of genes measured, though the magnitudes of expression differences were small. Coexpression patterns were more strongly preserved by race group for Oncotype DX and Decipher versus Prolaris (integrative correlations of 0.87, 0.73, and 0.62, respectively). Poorer prognosis was estimated in EAM versus AAM for Oncotype DX (p < 0.001), whereas no difference in prognosis was predicted between AAM and EAM using Prolaris or Decipher (p > 0.05). Replication of our findings directly on the commercial panels with long-term follow-up is warranted.

**Conclusions:** Due to observed racial differences across three commercial gene expression panels for PCa prognosis, caution is warranted when applying these panels in clinical decision-making in AAM.

## 1. Introduction

Prostate cancer (PCa) disproportionately affects African American men (AAM) compared to European American men (EAM), where AAM are nearly 70% more likely to be diagnosed with PCa and more than twice as likely to die from the disease compared to their EAM counterparts [1]. While differences in access to care and socioeconomic status are contributing factors to PCa disparities, accumulating evidence suggests the molecular genetic drivers implicated in carcinogenesis and PCa progression exert their effects in a race-specific manner [2]. In particular, genetic variants associated with PCa risk at the 8q24 and 17q21 loci are more common in AAM than EAM [3,4]. However, known identified risk SNPs appear to have limited clinical relevance regarding prognosis [5]. Rather, gene expression profiles from tumor tissue are more commonly used along with clinicopathologic features to predict risk of progression or fatal disease. Finally, although some studies have explored differences in gene expression patterns across race, the clinical implications of such differences remain unclear [6-8].

Current National Comprehensive Cancer Network (NCCN) guidelines recommend three gene expression-based tests for PCa prognosis in men with low or favorable intermediate risk disease: Decipher, Oncotype DX Prostate, and Prolaris [9]. Decipher, which was developed by GenomeDX, consists of a 22 marker panel covering 19 genes that produces a genomic risk score between 0 and 1 [10]. This test was developed on the Affymetrix Human Exon 1.0 ST array, and is recommended for predicting recurrence or metastases post-radical prostatectomy (RP) in patients with adverse pathology. Genomic Health created Oncotype DX Prostate to predict adverse pathology after RP for men with low to very low risk disease and with a 10 to 20 year life expectancy. The test, developed on the QRT-PCR platform, utilizes 12 genes involved in stromal response, cell morphology, androgen signaling and proliferation to calculate a Genomic Prostate Score that ranges from 0 to 100 [11]. The Prolaris test from Myriad Genetics was developed on the QRT-PCR platform, and calculates a risk score from 31 cell cycle progression (CCP) genes. Prolaris is recommended for post-biopsy RP and for untreated patients that have low or very low risk and a life expectancy of at least 10 years [12]. Notably, all three tests were developed in predominantly EAM cohorts, where Decipher, Oncotype DX Prostate, and Prolaris leveraged patient populations with 86%, 76%, and 94% EAM, respectively [10-13].

To date, no head-to-head comparisons of the three commercial tests have been conducted with long-term patient follow-up, though recent evidence suggests that the tests may provide inconsistent risk estimates in the presence of multifocal disease [14-15]. Limited data exist to explore how these genomic risk scores correlate with observed racial PCa disparities. In this study, we assessed if the genes that comprise the three prognostic gene signatures are differentially expressed by race in an independent genomic platform, and if general expression patterns are conserved between racial groups in a large cohort of EAM and AAM PCa patients.

## 2. Patients and methods

A total of 327 PCa patients contributed gene expression and clinical data to this study. These patients were selected from a cohort of 2,725 men with PCa treated with RP and bilateral pelvic lymph node dissection at the Moffitt Cancer Center (Tampa, FL), and which had Formalin-Fixed Paraffin Embedded (FFPE) tumor tissue available for research purposes. All AAM in the cohort with available tissue specimens were included (N = 95); the remaining 232 EAM cases were randomly selected based on tumor tissue availability. This study was approved by the Institutional Review Board at the Moffitt Cancer Center. Patient clinical data includes information on age, race, clinical stage, clinical Gleason on diagnostic biopsy, preoperative PSA levels, surgical pathologic information (tumor grade, stage, surgical margins status (SM), extraprostatic extension (ECE), or seminal vesicle involvement (SVI), and lymph node involvement (LVI)). Post-surgical risk status of patients was calculated using the Cancer of the Prostate Risk Assessment score (CAPRA-S), ranging from 0 - 12, and classified as low (0 - 2), intermediate (3 - 5) and high risk (6 - 12) [16]. Patients with missing preoperative PSA who did not have adverse pathologic features of prostatectomy *(i.e.,* SM, ECE, SVI and LNI) and had Gleason score ≤ 3+4 were considered low risk (CAPRA-S of 0 - 2). NCCN pretreatment risk was also categorized as low (PSA < 10 ng/mL, Gleason 3+3 and T stage < 2A), intermediate (PSA > 10-20 ng/mL or Gleason score = 7 or T stage was 2b-2c) and high (PSA > 20 ng/mL or Gleason score > 8 or T stage > 3b) [17].

All tumor samples included in this study passed stringent Nanostring gene expression quality control metrics. Gene expression was quantified for 60 genes across the three commercial panels using the Nanostring nSolver software. Eight house-keeping (HK) genes were used for normalization: *ATP5G3, EIF2B2, GAPDH, HMBS, MRPS9, PCBP2, RPA2* and *UBB.* Samples were excluded based on QC flags from the NanoString nSolver software. Additional exclusion based on the raw counts included: geometric mean for the HK gene counts < 100, the geometric mean for the HK genes < geometric mean for Negative control probes + 2 times their standard deviation. Additional samples were excluded based on principal component analyses and low expression of *KLK2* and *KLK3* (PSA, prostate-specific antigen) indicating lack of prostate cells in the sample.

Three genes are targeted by duplicate probesets on the Decipher panel *[AN07, MYBPC1, UBE2C,* and these genes were only placed once our Nanostring array. One gene, *CDCA8,* from the Prolaris panel, was not placed onto our array.

For each gene, expression was compared between EAM and AAM using Mann-Whitney tests, and log2 median fold changes were calculated to assess difference magnitudes. Spearman’s correlation was used to estimate gene co-expression within race, as well as the race-specific correlation between gene correlations, also known as the integrative correlation or the correlation of correlations method [18]. Partial Spearman’s correlations controlling for CAPRA-S group were also calculated. A continuous risk score for each gene panel was approximated using methods described elsewhere [14]. Confidence intervals for reported correlations were obtained by bootstrapping with 1,000 replicates.

All analyses were performed in R version 3.5.0. Data are available in Supplementary Table 1 and code and data to fully reproduce the analysis is available at https://github.com/GerkeLab/prostate PrognosticPanels.

## 3. Results

Clinical characteristics of the patients in the study cohort were broadly similar by race, with the exception that AAM patients had a lower median age at diagnosis than EAM patients (54 yrs *vs.* 60 yrs, p= 2.82E-6; Table 1). Approximately half of the genes measured across the gene panels 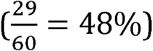 were differentially expressed by race using a Mann-Whitney threshold of *p* < 0.05 (Fig. 1). All 13 of 30 differentially expressed Prolaris genes showed higher expression in EAM compared to AAM. Among the 7 of 12 differentially expressed Oncotype DX genes, 4 were overexpressed in EAM and 3 were overexpressed in AAM. Finally, from the 19 Decipher genes, 8 were overexpressed in EAM and 1 was overexpressed in AAM. The proportion of genes overexpressed among AAM was highest in Oncotype, and lowest in Prolaris (43%, 11%, and 0% for Oncotype, Decipher, and Prolaris, respectively).

**Table 1:**
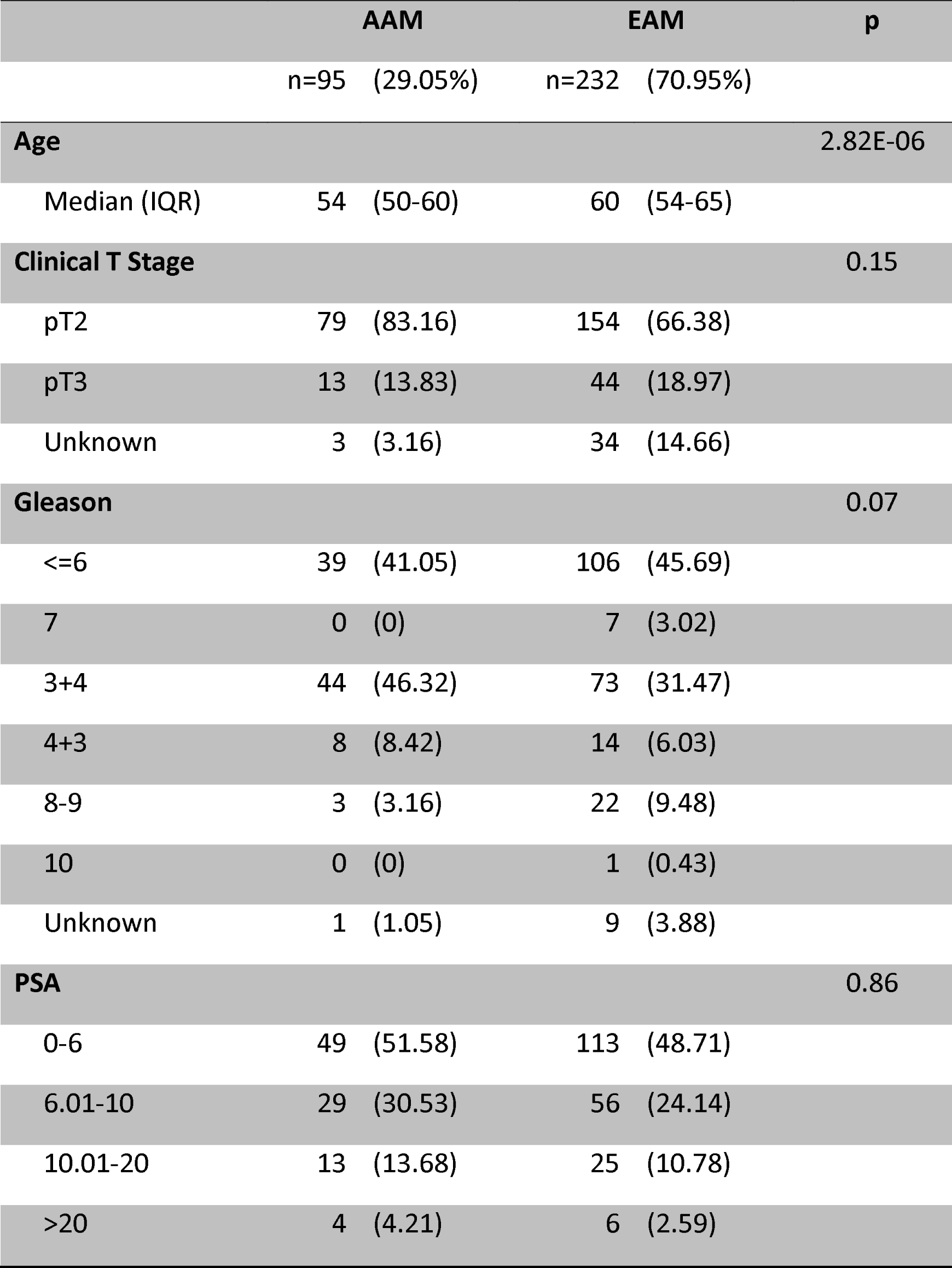
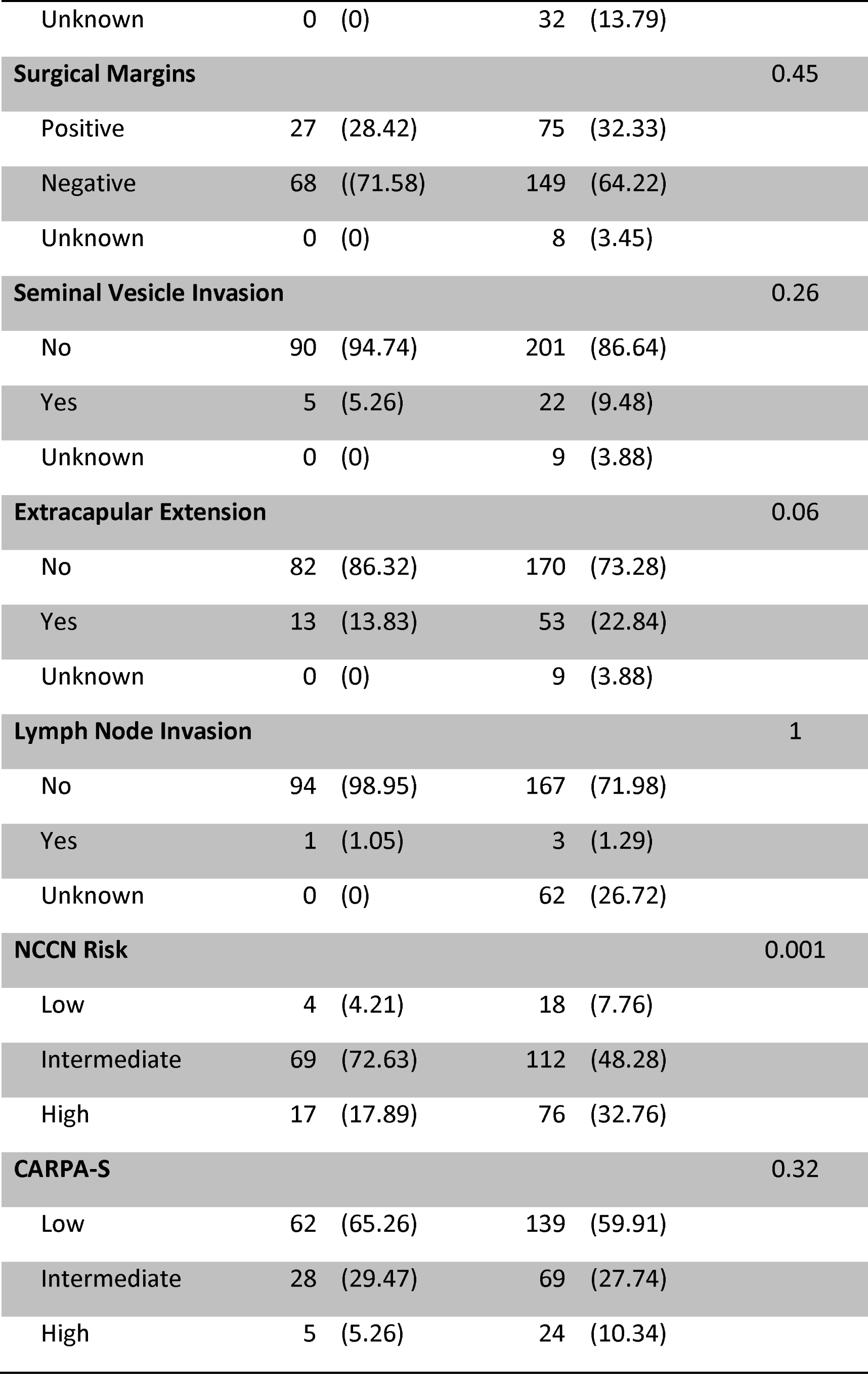
Baseline characteristics of the study population. *P* values were calculated using t-tests and Mann Whitney tests for continuous and categorical variables, respectively.

**Figure 1:**
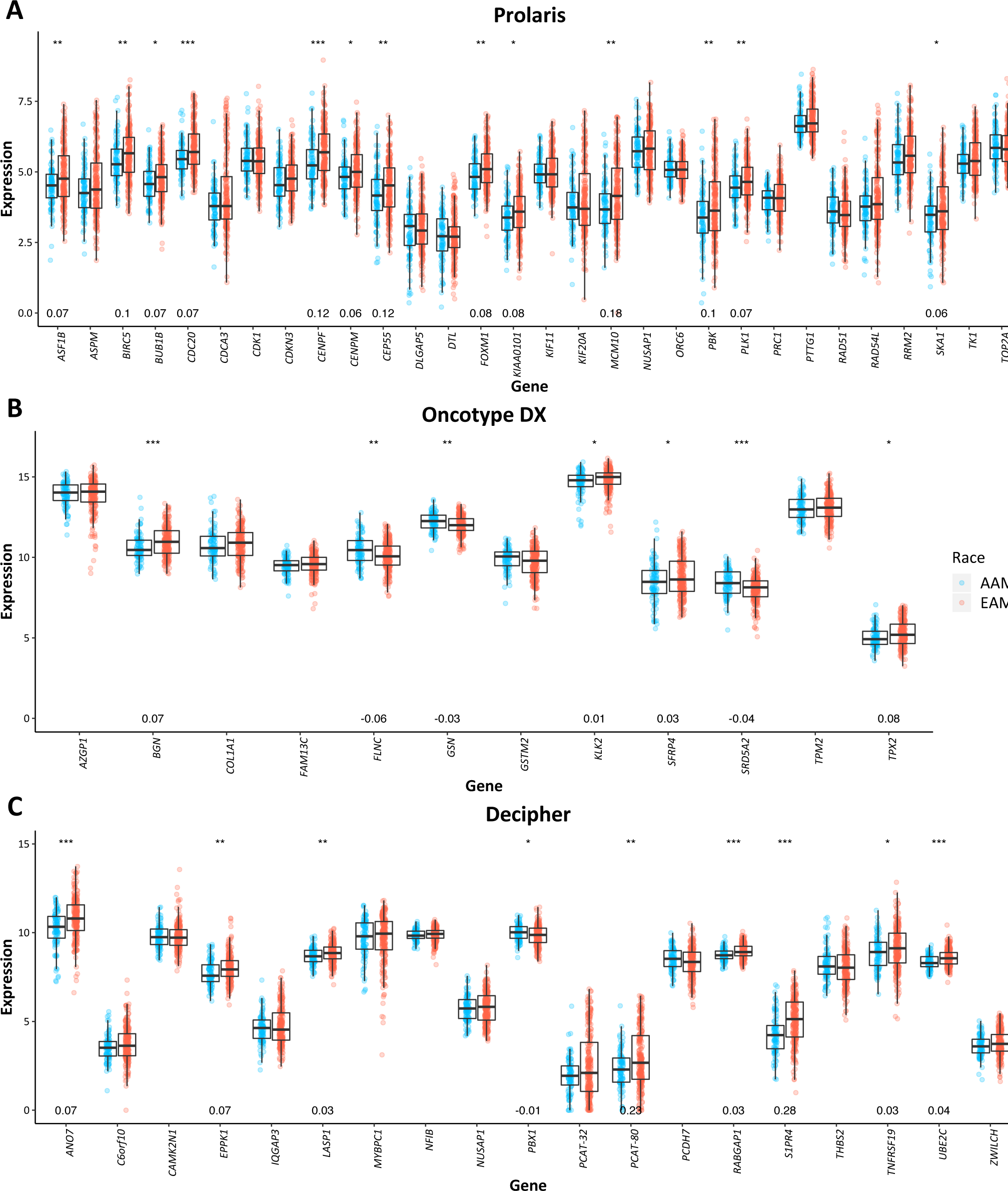
Gene expression for genes utilized by Prolaris, Oncotype and Decipher. *** denotes *p* < 0.001, ** *p* < 0.01, * *p* < 0.05. Log2 median FC is shown for those with a statistically significant difference in gene expression between EAM and AAM.

Within EAM, all Prolaris genes were positively correlated with one another, and this pattern was maintained in AAM but to a weaker extent (Fig. 2A). A moderate correlation in Prolaris gene correlations between EAM and AAM was also evident, 0.62 (0.56-0.69) (Fig. 2A), and this correlation was unchanged when adjusted for CAPRA-S, 0.62 (0.56-0.68). Little difference in co-expression was observed between EAM and AAM within Oncotype DX (Fig. 2B). Associations between intra-panel gene correlations and race were strong in Oncotype DX, 0.87 (0.81-0.97) (Fig. 2B), even when accounting for CAPRA-S, 0.88 (0.83-0.98). In the Decipher panel, gene correlations were more often positive and weak in AAM compared to EAM (Fig. 2C). The overall and CAPRA-S weighted correlation of correlations were similar, with 0.73 (0.66-0.82) for the unweighted and 0.74 (0.68-0.82) for the CAPRA-S weighted.

**Figure 2:**
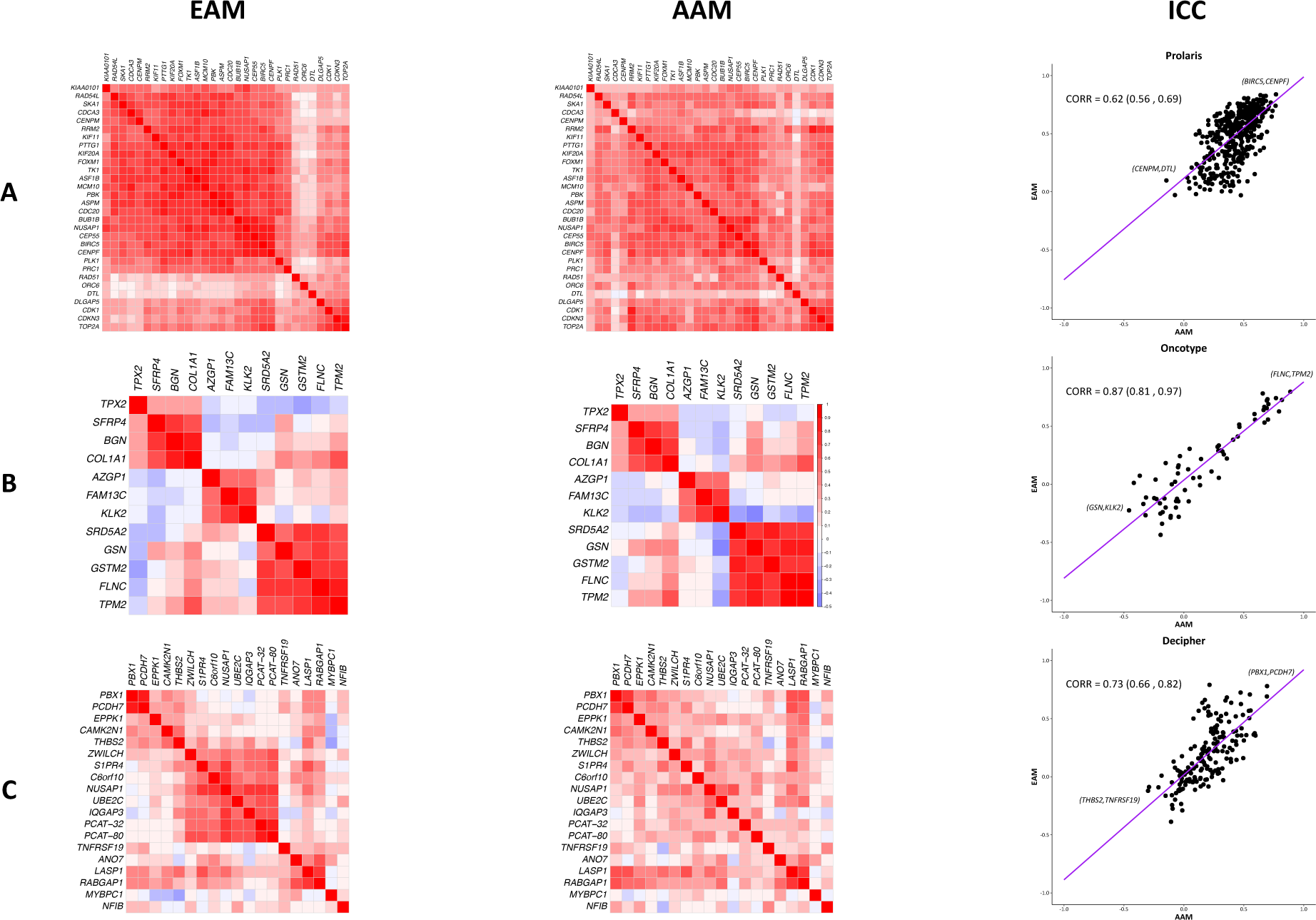
The panels are Prolaris (A), Oncotype Dx (B) and Decipher (C). Heatmap of inter-gene Spearman’s correlations, EAM (left) and AAM (right), are shown in the first two columns. CAPRA-S weighted integrative correlation coefficients and 95% confidence intervals for genes in each panel are shown in the far right column. Gene pairs with the highest and lowest correlation in each panel are annotated.

Surprisingly, the estimated genomic risk scores predicted more adverse outcomes in EAM compared to AAM in Oncotype DX (p=0.0004), whereas estimated genomic risk predicted similar outcome for AAM compared to EAM when using Prolaris (p=0.21) and Decipher (p=0.29) genes (Fig. 3). When comparing genomic risk scores between panels (Fig. 4), consistent, moderately positive correlations were observed for both EAM and AAM in all pairings. Decipher risk scores were positively correlated (p<0.05) with CAPRA-S and NCCN risk group in EAM but insignificantly correlated in AAM. Prolaris risk was not correlated with either clinical score in EAM or AAM. Oncotype was positively correlated with CAPRA-S in EAM and NCCN risk groups in both EAM and AAM.

**Figure 3:**
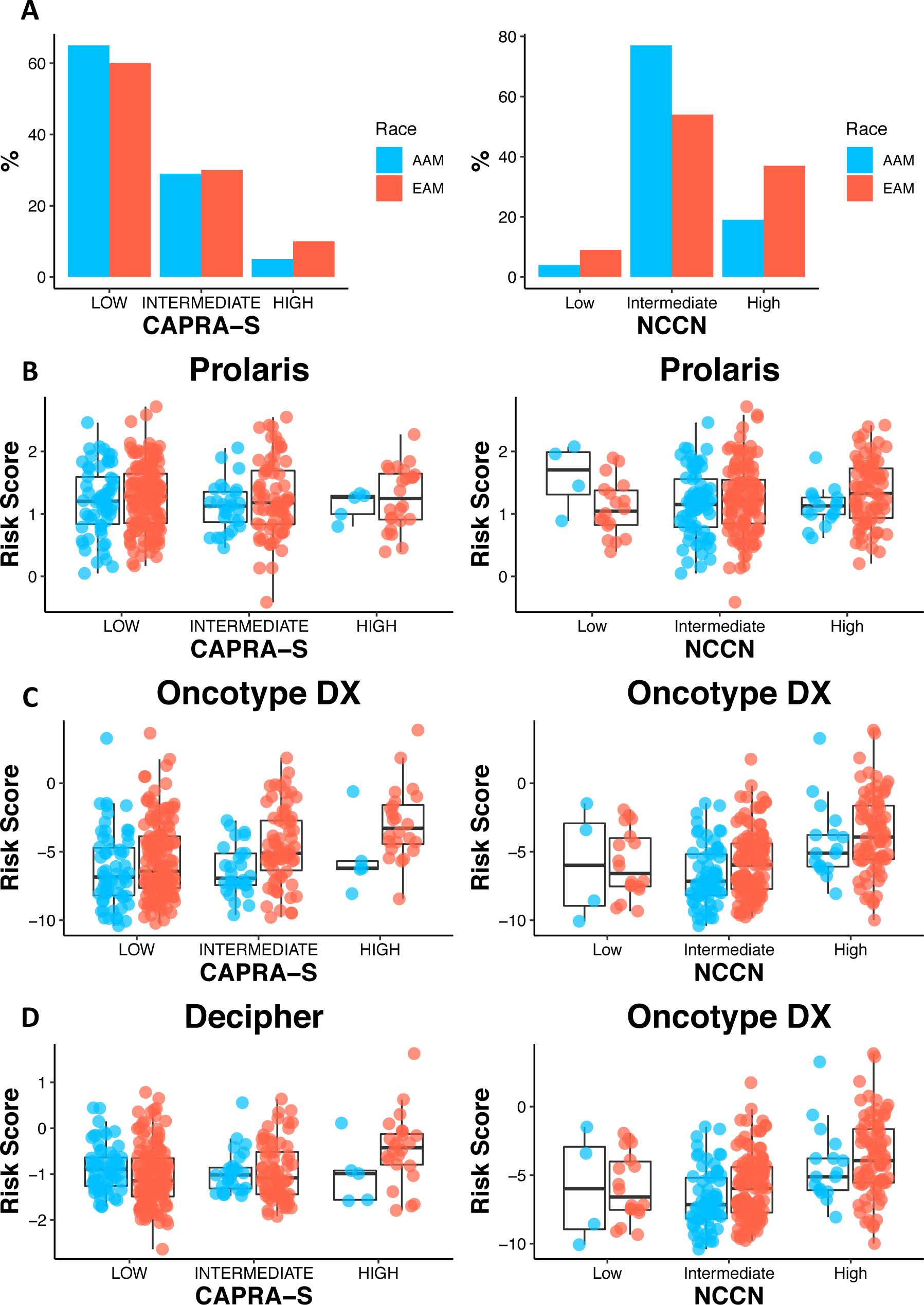
Percentage of EAM and AAM patients with low (0-2), intermediate (3-5), and high (6-12) CAPRA-S and with low, intermediate and high NCCN risk classification. Overall risk scores for Prolaris (B), Oncotype Dx (C) and Decipher (D) within CAPRA-S and NCCN groups.

**Figure 4:**
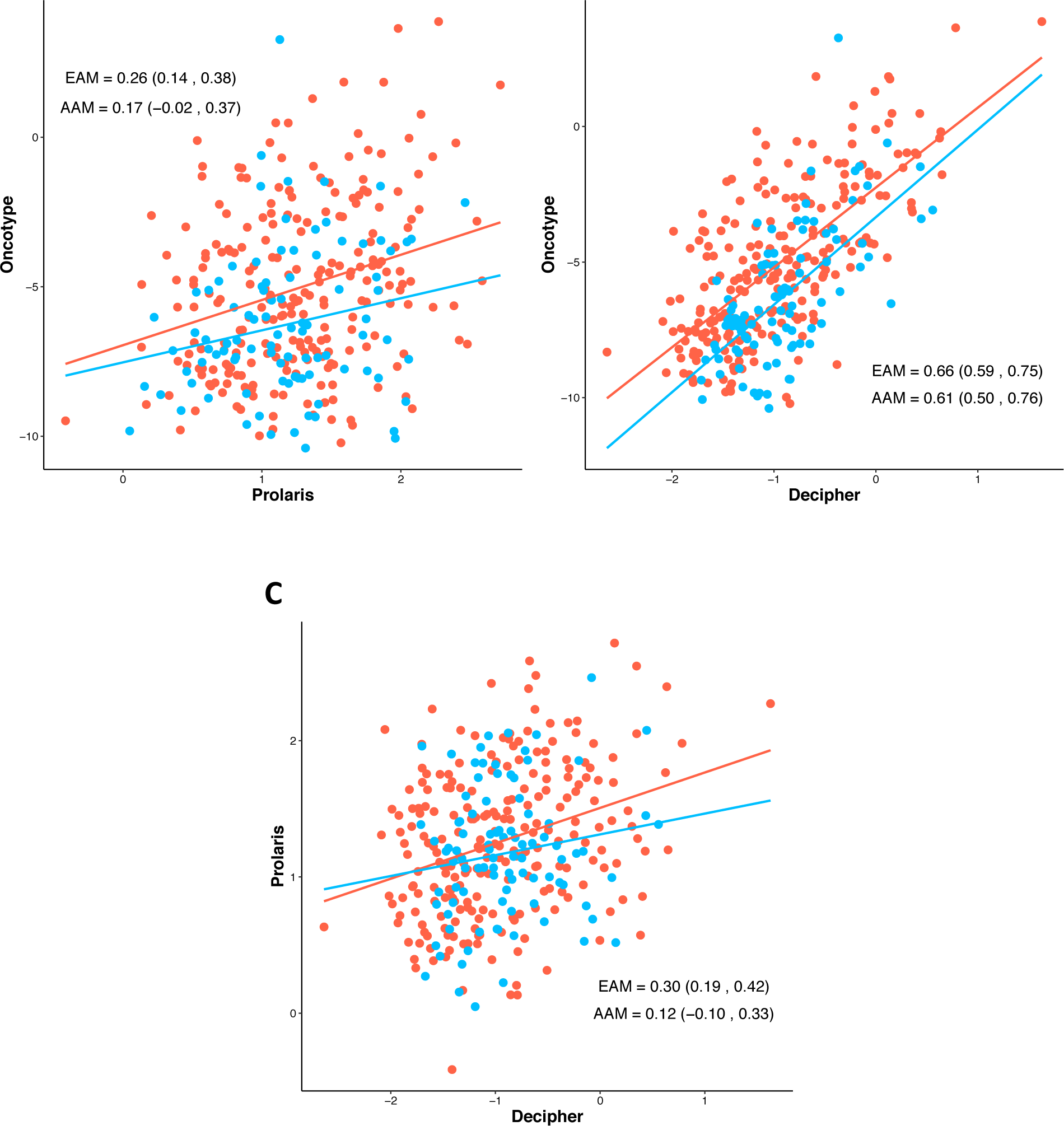
Spearman’s correlation and 95% confidence intervals of overall risk scores between each panel and within each race.

## 4. Discussion

We examined differences in expression between AAM and EAM among genes included in three commercially available PCa prognostic biomarker panels: Prolaris, Oncotype DX, and Decipher. Broadly, we hypothesized that racial differences in expression patterns across the panels would exist, given the evidence that both underlying biology and predicted prognoses significantly differ by race.

In general, significant racial differences were observed for individual genes and for correlational patterns within panels, albeit relatively small in magnitude. Of the 60 genes examined, 48% were differentially expressed between EAM and AAM, and the panel with the highest proportion of differentially expressed genes was Oncotype DX (58%), followed by Decipher (47%) and Prolaris (43%). However, while nearly half of the genes were differentially expressed, the magnitude of median expression differences was relatively small, with the greatest being *S1PR4* (encoding Sphingosine-1-Phosphate Receptor-4) in Decipher where there was 28% higher median expression in EAM. The range of differences we observed was consistent with that of genes found to be differentially expressed by race in other studies [2]. Inter-gene correlational patterns were generally preserved within each of the three panels, although with somewhat weaker signals in AAM patients.

Inconsistent with current clinical data indicating that AAM tend to experience less favorable outcomes than EAM, the Decipher and Prolaris estimated genomic risk scores showed no worse prognosis in AAM compared to EAM. Paradoxically, we found that better prognosis was estimated by Oncotype DX genes for AAM compared to EAM patients. For each patient, the risk score estimates across the three panels showed a somewhat weak positive correlation within racial groups.

To our knowledge, this is the first analysis evaluating race-specific expression patterns of established prognostic genes in three commercially available panels on a single gene expression platform. Results from this study provide evidence for caution when applying genomic predictors developed in predominantly EAM to AAM with PCa, and emphasizes the importance of conducting de novo genomic studies in samples derived from at-risk AAM populations. However, this study is not without limitations. In interpreting our findings with respect to risk scores, it is important to note that we have quantified gene expression by Nanostring, rather than by the respective commercial panels, and we therefore needed to approximate the relative weights given to each gene in the signature. Nonetheless, we were able to leverage directional associations of the genes with outcomes from the original studies, and our approach to sum the signed expression values has been proven successful in other contexts [14-15,19].

Additionally, due to the lack of long-term follow up data in our cohort the implications of our study for the use of these panels in AAM remains to be determined. Specifically, without extended follow-up, we are at this time unable to provide direct evidence that any one test more robustly captures a signal that is relevant to racially disparate outcomes in PCa. However, our findings do clearly indicate that further research is needed to determine if prognostic gene expression signatures should be applied in a race-sensitive manner.

## 5. Conclusions

Although prognostic accuracy of commercially available gene expression tests for PCa has been consistently demonstrated in EAM, our findings clearly indicate that a thorough assessment of the performance of these panels in AAM is needed. Our study identified several gene-specific differences and correlational patterns comparing EAM to AAM, though these differences were low in magnitude. Interestingly, none of the approximated risk scores indicated a markedly higher risk for AAM compared to EAM, a result that is counter-intuitive with respect to broader clinical trends. Future studies that validate the prognostic value of such tests in AAM are needed.

## Data Availability

https://github.com/GerkeLab/prostatePrognosticPanels

## Author Contributions

Jordan Creed and Travis Gerke have full access to all the data in the study and take responsibility for the integrity of the data and the accuracy of the data analysis.

*Study concept and design:*Creed, Gerke, Yamoah

*Acquisition of data:*Creed, Berglund, Awasthi, Rounbehler

*Analysis and interpretation of data:*Creed, Berglund, Gerke

*Drafting of the manuscript:*Creed, Yamoah, Cleveland, Gerke

*Critical revision of the manuscript for important intellectual*

*content* Creed, Gerke, Berglund, Park, Cleveland, Yamoah

*Statistical analysis:*Creed, Gerke, Berglund

*Obtaining funding:*Cleveland, Park, Yamoah, Berglund, Gerke

*Administrative, technical or material support:* Awasthi, Berglund, Rounbehler

*Supervision:*Gerke

*Other:*None

## Informed Consent

The appropriate Institutional Review Board approval was obtained for the study protocol at Moffitt Cancer Center (IRB protocol 105514).

## Acknowledgements

This work is supported in part by the Molecular Genomics Core Facilities at the Moffitt Cancer Center through its NCI CCSG grant (P30-CA76292). The authors would like to thank Brandon Manley and Julio PowSang for their valuable input on this manuscript. We sincerely thank the participants of this study for their contributions.

